# Imaging translation dynamics in live embryos reveals spatial heterogeneities

**DOI:** 10.1101/2020.04.29.058974

**Authors:** Jeremy Dufourt, Maelle Bellec, Antonio Trullo, Matthieu Dejean, Sylvain De Rossi, Mounia Lagha

## Abstract

The translation of individual mRNA molecules is a key biological process, yet this multi-step process has never been imaged in living multicellular organisms. Here we deploy the recently developed Suntag method to visualize and quantify translation dynamics of single mRNAs in living *Drosophila* embryos. By focusing on the translation of the conserved major epithelial-mesenchymal transition (EMT)-inducing transcription factor Twist, we identified spatial heterogeneity in mRNA translation efficiency and reveal the existence of translation factories, where clustered mRNAs are co-translated preferentially at basal perinuclear regions. Simultaneous visualization of transcription and translation dynamics in a living multicellular organism opens exciting new avenues for understanding of gene regulation during development.

## Introduction

During the development of multicellular organisms, gene expression must be precisely regulated in time and space. Decades of genetic manipulations in *Drosophila* have dissected the gene regulatory networks responsible for the establishment of precise patterns of gene expression(*1*). The establishment of these patterns has been primarily studied at the transcriptional/mRNA level in fixed embryos using fluorescent in situ hybridization (FISH), or in living embryos via the recent adoption of mRNA labeling technologies such as the MS2/MCP system(*2-4*). In the context of a developing embryo, precision in mRNA production is functionally relevant only if it leads to precision in protein abundance.

While this central dogma has been established over the last 60 years, the non-linearity between the levels of a given mRNA and the amount of protein it encodes has only recently started to emerge(*5-8*). One explanation for this poor correlation resides in the spatial control of translation that could lead to differential protein output. It is indeed well established that mRNAs can be localized to particular subcellular compartments(*9*). This subcellular targeting is evolutionary conserved and has been described in a variety of organisms from bacteria to mammals(*10*). However, the consequences of such localization in terms of translational output are much less thoroughly understood. In some cases, localization of mRNAs favors their translation, as exemplified by *oskar* at the posterior pole of *Drosophila* oocytes(*11*) or β-actin at neuronal branching points(*12, 13*). But in other cases, mRNAs can assemble in multimolecular complexes to prevent their translation, such as with mRNAs located in stress granules(*14*).

The non-linearity between mRNA and protein distributions has primarily been studied with genomic approaches on populations of cells where spatial constraints are eroded. Indeed, while live imaging of mRNA has been possible since 1998(*15*), a similar method to image many cycles of translation (Suntag) was only deployed in 2016 in cultured cells(*13, 16-19*), and has yet to be introduced in an intact developing organism. Here we have implemented the Suntag labeling method in *Drosophila* and quantified the translational dynamics of endogenous *twist* (*twi*) mRNA molecules in living embryos. We uncovered a broad range of translation efficiencies of the *twi* mRNA population. In particular, cytoplasm located below nuclei fosters enhanced translation, in part via directed-mRNA clustering in translation factories. The revealed translation factories of *twi* mRNAs are primarily located in the basal perinuclear space and exhibit reduced mobilities. Moreover we identified that single mRNA molecules located in this subcellular compartment are more efficiently translated than identical mRNA located apically. Localized and enhanced translation of transcription factor encoding mRNAs at the basal perinuclear cytoplasm may facilitate their nuclear import and limit their diffusion, two essential features allowing precise formation of primordial embryonic germ layers.

## Results

### Implementation of Suntag method to visualize translation in a developing organism

In the Suntag system, the multimerization of a GCN4-based epitope (named suntag), with high affinity for a single chain antibody scFv coupled to a fluorescent protein (sfGFP)(*20*) allows multiple scFv-GFP to bind (Figure 1a). This signal amplification creates a bright fluorescent spot above the background signal. We focused our analysis on the *twi* gene, which encodes a key conserved transcriptional activator of the mesodermal gene network and the EMT program in metazoans that is frequently deregulated in a large number of metastatic cancers(*21*).

**Figure 1:**
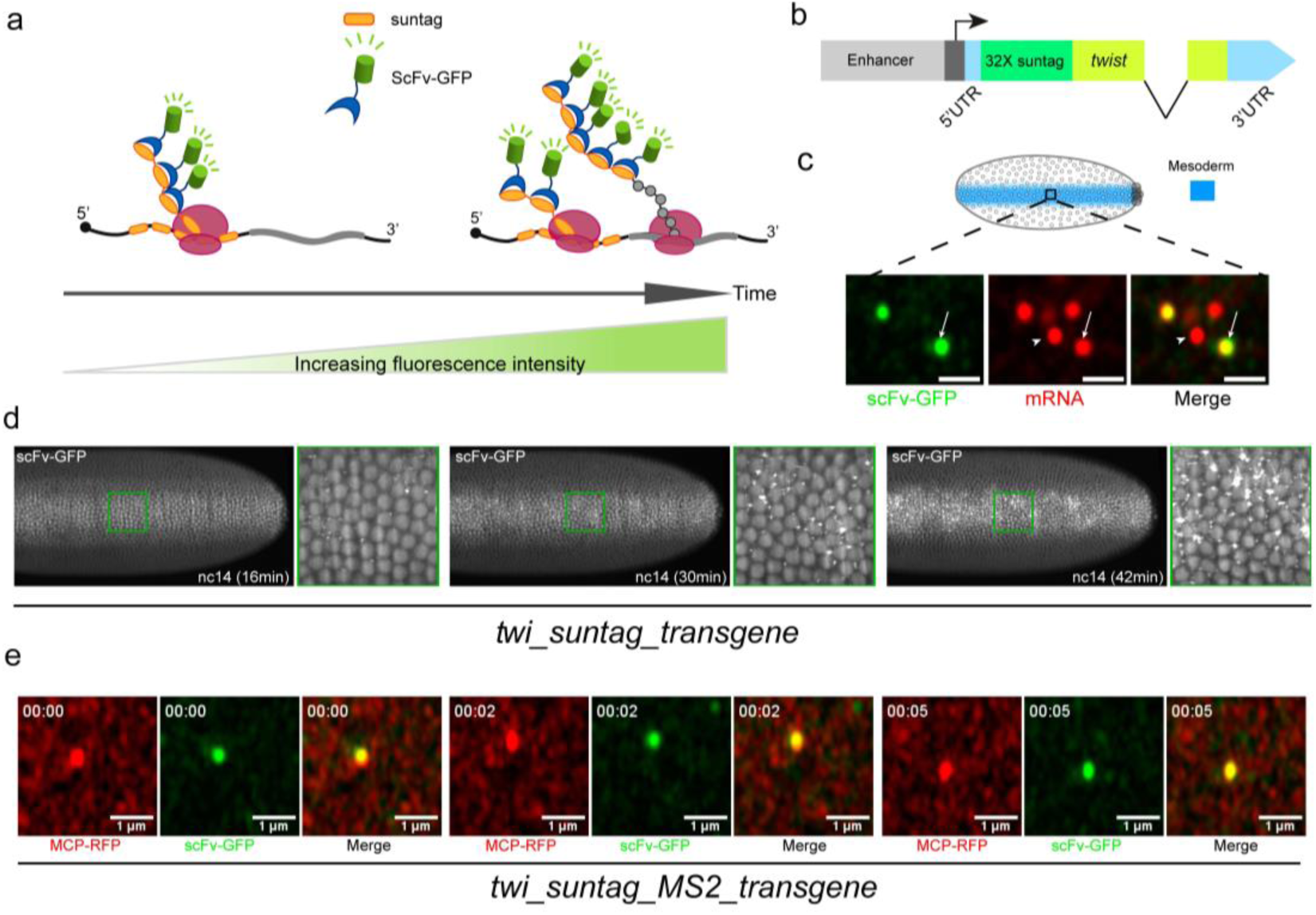
Imaging translation with suntag labeling in *Drosophila* embryos. **a**, Principle of the Suntag system: labeled antibody probes, here single-chain variable fragments coupled to sfGFP (scFv-GFP), bind to repeated suntag epitopes, allowing visualization of nascent translation. Tandem array of suntag peptides are shown in orange, scFv-GFP molecules in green and ribosomes in purple. **b**, Schematic representation of *twist_Suntag* transgene. **c**, Zoomed confocal single plane within the mesoderm of a *scFv-GFP-NLS/+ > twi_suntag/+* embryo stained with endogenous scFv-GFP (green) and probes against suntags (red). scFv-GFP foci colocalizing with suntag probes reveal mRNAs in translation (yellow, arrow), while spots that are only stained with suntag probes correspond to non-translating mRNAs (red, arrowhead), scale bars 1μm. **d**, Snapshots of dorsal and ventral views of a maximum intensity projected Z-stack from a *scFv-GFP-NLS/+ > twi_suntag/+* live embryo imaged with a MultiView Selective Plane Illumination Microscope, MuViSPIM, (related movie 2). Associated zoomed images are shown on the right. All embryos are ventral views, oriented with anterior to the left. **e**, Frames taken from a fast-confocal movie (≈2 frames/sec) of a *scFv-GFP-NLS/MCP-TagRFPT-NLS > twi_suntag_MS2/+* embryo (related to movie 4), showing an mRNA molecule (red) co-localizing and moving with a bright scFv-GFP signal (green).

Initially, to test if the Suntag labeling system could be functional in a living organism we created a *twi_suntag* transgene whereby 32 suntag repeats(*18*) were inserted immediately after the start codon (Figure 1b). In addition, we created various scFv-GFP lines for maternal deposition, in embryos, of the scFv detector protein (Figure S1a-c and movie 1and S1). To ensure minimal levels of free scFv in the cytoplasm, we added a nuclear localization signal (NLS; Figure S1c and movie 1), referred to as scFv-GFP. We also generated a graded scFv-GFP line with high expression at the anterior pole that diminishes towards the posterior region (*scFv-GFP-bcd 3’UTR*) (Figure S1d). A diffuse, soluble GFP signal is detected in all scFv lines, showing that scFv can be genetically encoded in *Drosophila* without forming aggregates (Figure S1 b, c and movie 1 and S1).

In the presence of the *twi_suntag* transgene and scFv-GFP detector protein, clear fluorescent spots are detected in fixed embryos at nuclear cycle (nc) 14 specifically in the mesoderm, where *twist* is transcribed (Figure S1e).

While the suntag signal was restricted specifically to the mesoderm, *twi_suntag* expression appeared stochastically in cells within this domain, likely due to the presence of only the *twi* proximal enhancer in the transgene. Thus, our *twi* transgene did not fully recapitulate *twi* canonical expression (Figure S1f). By performing single molecule mRNA labeling using single molecule fluorescent *in situ* hybridization (smFISH) with the simultaneous detection of native scFv-GFP, we could detect two populations of cytoplasmic single mRNA molecules: those co-localizing with a bright GFP signal, ≈56% of total mRNA in nc14 (n=6231 from three smFISH), corresponding to mRNAs in translation, and those devoid of a GFP signal and corresponding to untranslated mRNAs (Figure 1c). By live imaging, distinct spots were detected clearly above background within the presumptive mesoderm of *twi_suntag* transgenic embryos (Figure 1d and movie 2). These bright spots correspond to nascent sites of translation, as they disappear upon puromycin injection into live embryos (Figure S1g and movie 3).

In order to visualize both mRNA and their translation in living embryos, we created another transgene where, in addition to the suntag sequences, MS2 repeats were inserted in the 3’UTR region (Figure S1h). To amplify the fluorescent signal produced by single molecules of mRNA, we employed a 128-loop array of a new generation of optimized MS2 loops(*22*). Similar to the *twi_suntag* transgene, *twi_suntag_MS2* transcripts were expressed in the presumptive mesoderm (Figure S1i) and we could detect two populations of cytoplasmic mRNAs, the first corresponding to mRNAs in translation (arrow) and the second to mRNAs not in translation (arrowhead) (Figure S1j).

Using two-color live imaging of the *twi_suntag_MS2* transgene (with scFv-GFP and MCP-TagRFPT detector transgenes), we occasionally detected mRNA spots co-localizing and traveling with suntag signal in the cytoplasm of living embryos (Figure 1e and movie 4). However, fast diffusion of mRNA molecules and rapid photobleaching of the fluorescent tag precluded their tracking. In conclusion, we have developed and validated the genetic tools to image translation in intact live *Drosophila* embryos. We can thus image translation in live embryos with scFv-GFP and use smFISH to detect and quantify single molecules of cytoplasmic mRNAs.

### Imaging translation from an endogenous locus

To monitor *twi* translational dynamics from the endogenous locus, we created a *twi_suntag_CRISPR* allele (Figure 2a) and a *twi_suntag_MS2_CRISPR* allele (Figure S2a). With its complete set of regulatory regions, *twi_suntag_CRISPR*and *twi_suntag_MS2_CRISPR* were activated throughout the mesoderm (Figure 2b, Figure S2b). The scFv-GFP protein labeling completely overlapped that of *twi* mRNAs labelled with smFISH probes directed against the suntag (Figure 2b) or MS2 sequences (Figure S2b), and bright scFv-GFP spots are observed in nc14. At single molecule resolution, bright foci of translating individual *twi* mRNA are detected in ≈64% of total pool of single mRNA molecules in nc14 (n=12022 from four smFISH), along with untranslated single mRNAs and single molecules of Twi protein (Figure 2c, Figure S2c). Using live imaging of these *twi_CRISPR* alleles, we examined the timing of *twi* translation. To image the entire embryo without compromising on the temporal resolution, we employed light sheet microscopy (MuViSPIM)(*23, 24*). This revealed that *twi* translation was strongly induced during nc14 (Figure 2d, 2e, Figure S2d, movie 5 and S2). This result was confirmed by smFISH and confocal microscopy (Figure S2e-f).

**Figure2:**
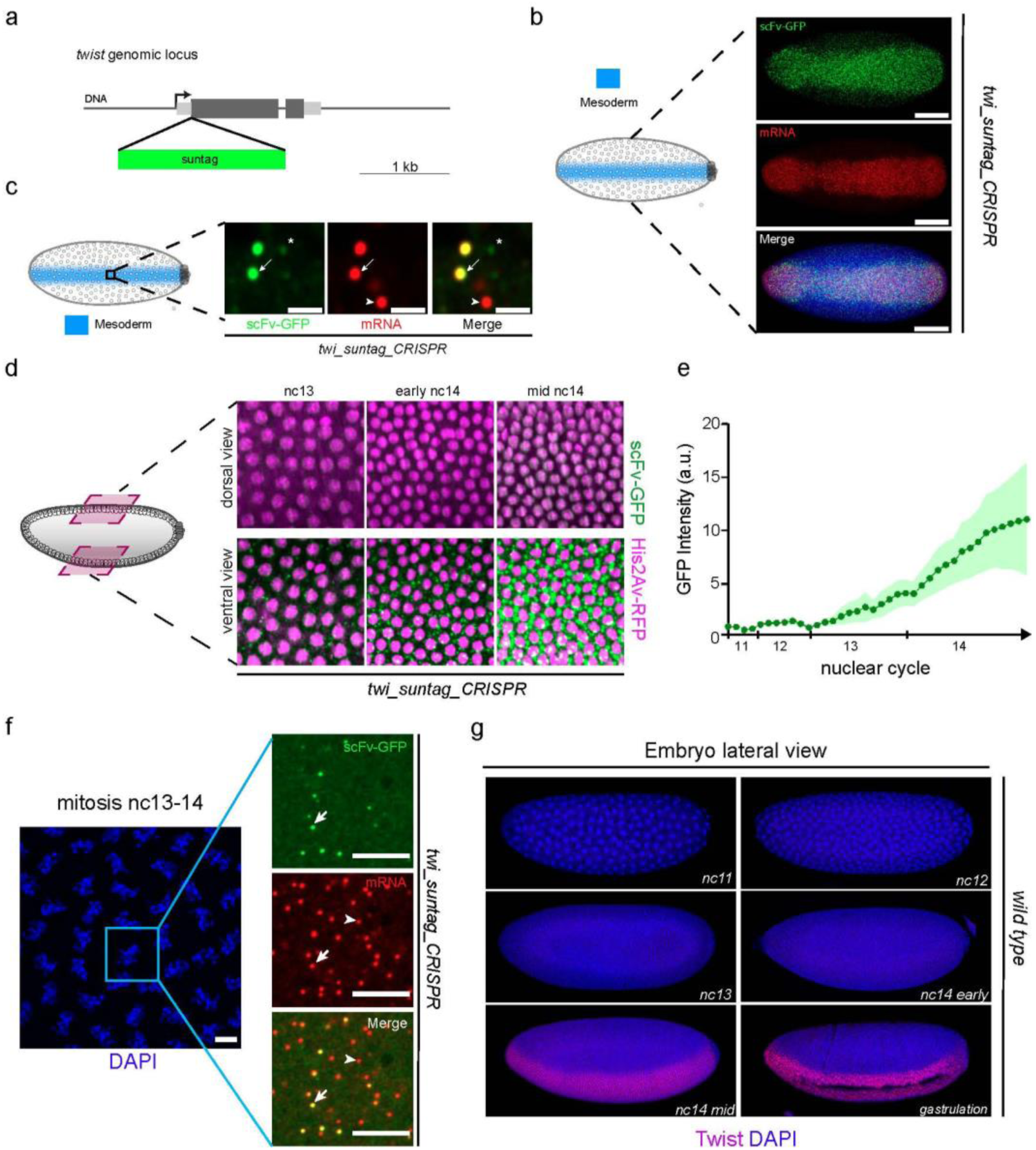
Capturing the timing of translation of endogenous *twist* mRNAs. **a**, CRISPR editing of endogenous *twist* gene by suntag repeats. **b, c**, Confocal images of nc14 *twi_suntag_CRISPR Drosophila* embryos expressing scFv-GFP (green) stained with a suntag smFISH probe (red). In **b**, Maximum intensity projection of a ventral view shows the *twi_suntag* is expressed within the entire *twi* pattern, scale bars 100μm. In **c**, zoomed view within the mesoderm exhibiting two groups of mRNAs (red) molecules: those co-localizing with scFv-GFP signal that are engaged in translation (arrows, yellow in merge) and those not co-localizing with a GFP signal that are untranslated (arrowheads, red only). Single molecule proteins are labeled only by scFv-GFP (stars). Scale bars 1μm. **d**, Dorsal and ventral maximum intensity projection snapshots of *His2Av-mRFP/+; scFv-GFP-NLS/+ > twi_suntag_CRISPR/+*, of a live embryo imaged with a MuViSPIM from nc13 to mid nc14 (Movie 5). Nuclei are shown in magenta and sites of translation in green. **e**, quantification of the scFv-GFP signal in the mesoderm from nc11 to 30 minutes into nc14, from which the ‘free’ scFv-GFP (unbound to suntag) measured in the dorsal side where *twi* is not expressed, was subtracted (n=3 embryos). **f**, Maximum intensity projection of three z-planes (≈1 μm) of confocal images of mitosis from nc13 to nc14 *scFv-GFP-NLS/+ > twi_suntag_CRISPR/+* embryos expressing scFv-GFP, labelled with suntag smFISH probes and DAPI, (left panel) large view of DAPI staining (blue). Right panels are zoomed views exhibiting two groups of mRNAs (red) molecules: those co-localizing with scFv-GFP signal (arrows, yellow in merge) and those not co-localizing with a GFP signal (arrowheads, red only), scale bars 5μm. **g**, wild type fixed embryos at the indicated developmental stage, stained with a Twi antibody (red) and DAPI (blue). All embryos are oriented with anterior to the left and ventral on the bottom.

Bright but rare scFv-GFP foci appeared as early as nc12 (Figure S2e-f) and we observe a persistent translation during mitoses (Figure 2f). Following the initial syncytial development stage, the plasma membrane progressively invaginates between adjacent nuclei during nc14 with an apico-basal directionality, a process named cellularization. Interestingly, the large wave of twi translation during nc14 occurs prior to the completion of cellularization. Thus, *twi* mRNA are efficiently translated during a developmental window where short range diffusion between neighboring ‘pseudo-cells’ is possible(*25*). The timing of *twi* translation is consistent with dynamics of *twi* mRNA production. *twist* is a zygotic gene which is among the first genes to be transcribed (∼nc11)(*26*). The number of cytoplasmic *twi* transcripts peaks early in nc14(*26*) and seems to precede the stage when *twi* translation is at maximum (Figure 2e).Moreover, the timing of *twi* translation is consistent with the timing of the appearance of nuclear Twi protein (Figure 2g). Given the pivotal role of Twi protein acting as a major transcriptional activator of the mesodermal fate(*27*), this relatively late translation was not expected.

By combining suntag and MS2 labeling, it is possible to image transcription and translation from an endogenous gene within a single embryo (Figure S2g and movie S3). Thus, double labeling with suntag/MS2 repeats provides a method to quantify the precise chronology of events and possible flow of information from transcription to translation.

### Spatial heterogeneity of twist translation

Having determined the precise timing of *twi* translation, we then investigated its subcellular spatial localization. Using are constructed transverse view of a developing embryo by MuViSPIM, the sites of translation in nc14 appeared much more prominent in the basal perinuclear region (i.e. towards the inside of the embryo),although translation was also observed in the apical perinuclear region (Figure 3a and Figure S3a and movie 6 and S4). To confirm the observation that translation is heterogeneously distributed, we quantified the scFv-GFP signal in these two compartments in living embryos from nc11 to nc14 by confocal microscopy (Figure S3b and S3c). Contrary to early developmental stages, where translation seems equivalent in the apical and basal cytoplasmic space, there is a striking difference in the behavior in nc14. In nc14, the largest and brightest spots of *twi* translation appeared mainly in the cytoplasm located below the nuclei (basal) (Figure 3a, S3a-c). To better characterize these large scFv-GFP foci with their associated mRNA corresponding to messenger ribonucleoproteins (mRNPs),we systematically quantified the number of translated and non-translated mRNAs in fixed samples in 3D (Figure 3b and S3d). While mRNA molecules were present along the entire depth of a cell volume (Figure 3c), their intensity was clearly enhanced at the level of the basal perinuclear space (Figure 3c-d). A similar spatial heterogeneity is observed for the scFv-GFP translation signal, which peaks in the basal perinuclear space (Figure 3c-d). We then determined the intensity of a single mRNA molecule from smFISH experiments and identified clusters of multiple molecules of mRNAs (see methods). These mRNA clusters were of varying sizes ranging from 2 to 6 mRNAs and were significantly more frequent in the basal perinuclear cytoplasm (Figure 3e).

**Figure3:**
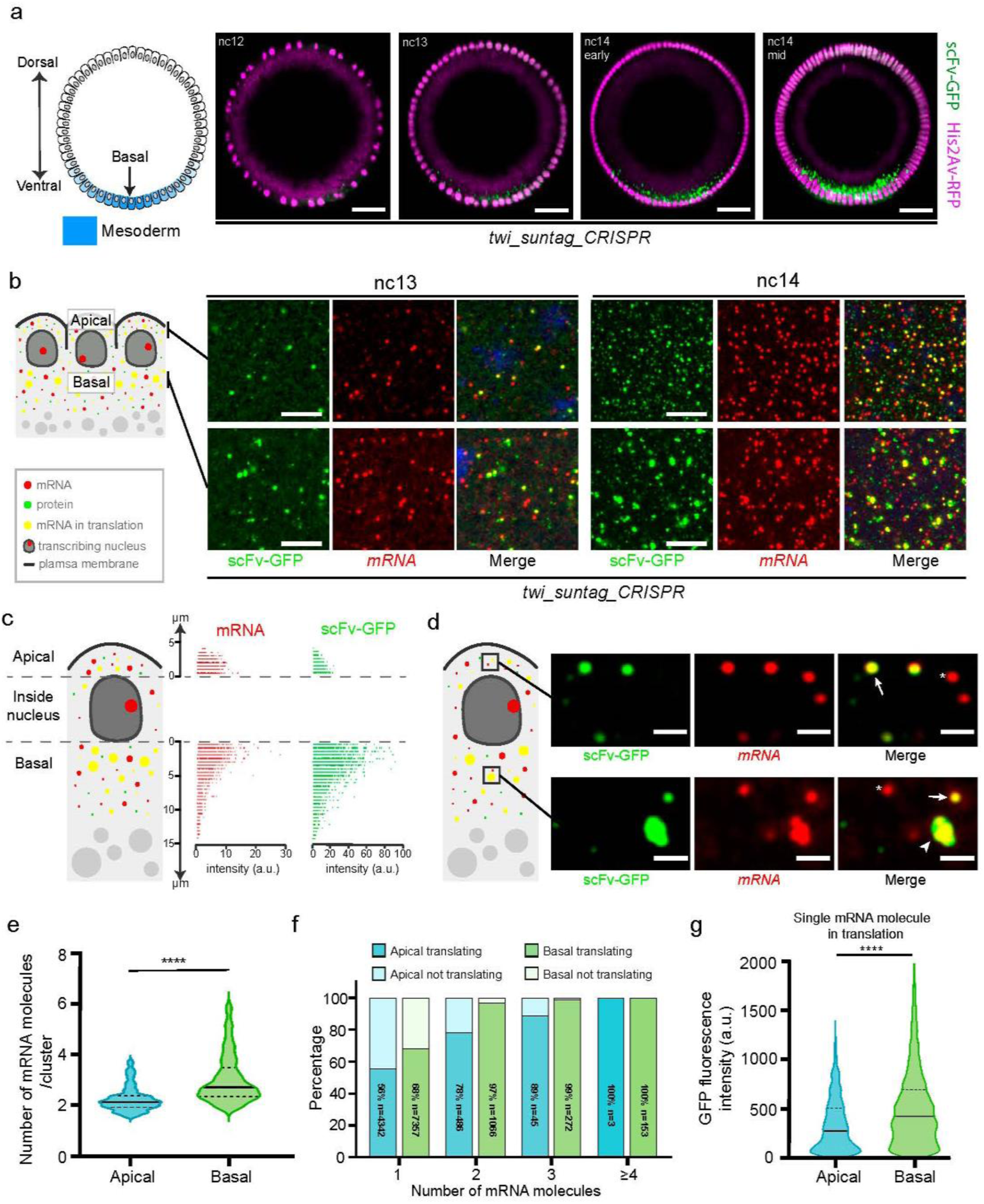
Revealing a spatial heterogeneity of endogenous *twist* mRNAs translation. **a**, Cross section of a transverse plane from reconstructed images by MuViSPIM of a *His2Av-mRFP/+; scFv-GFP-NLS/+ > twi_suntag_CRISPR/+* embryo (related movie 6). Ventral mesodermal nuclei are located at the bottom. Nuclei are observed with the His2Av-mRFP (magenta) and sites of translation with the scFv-GFP (green dots), scale bars 30μm. **b**, Maximum intensity projection of three z-planes (≈1 μm) of confocal images of nc13 and nc14 *scFv-GFP-NLS/+ > twi_suntag_CRISPR/+* embryos expressing scFv-GFP (green) labelled with suntag smFISH probes (red) and DAPI (blue), scale bars 5μm. **c**,Intensity quantification of detected mRNA spots (red) and scFv-GFP spots (greenin z-planes (0.5 μm apart) of four nc14 *scFv-GFP-NLS/+ > twi_suntag_CRISPR/+* embryos expressing scFv-GFP labelled with suntag smFISH probes. **d**, Zoomed z-projected confocal images taken from the apical and basal perinuclear compartment of *scFv-GFP-NLS/+ > twi_suntag_CRISPR/+ Drosophila* embryos expressing scFv-GFP (green) and-labelled with suntag smFISH probes (red). Top panels show two z-projected images of the apical compartments. Bottom panels are equivalent z-projected images located basally on which it is possible to distinguish single mRNA molecules not in translation (white stars), single mRNA molecule engaged in translation (white arrows), and large GFP foci colocalizing with cluster of mRNAs (arrowhead). **e**, Violin plot of the distribution of the number of mRNA molecules per cluster from images taken apically (n=523; blue) and basally (n=1384; green) from four nc14 embryos *scFv-GFP-NLS/+ > twi_suntag_CRISPR/+* labelled with *suntag* smFISH probes. Centered black bar represents the median, dashed black lines represent quartiles. ∗∗∗∗ p < 0.0001 with a two-tailed Welch’s t-test. **f**, Percentage of mRNA in translation located in the apical or basal cytoplasm(blue or green respectively) or not translating (light blue or light green respectively) in each class of mRNA pool (see main text and methods), from four *scFv-GFP-NLS/+ > twi_suntag_CRISPR/+* nc14 embryos labelled with suntag smFISH probes.**g**, Violin plot of the distribution of scFv-GFP intensities colocalizing with single mRNA molecules located apically (n=2380; blue) and basally (n=4202; green) from four nc14 *scFv-GFP-NLS/+ > twi_suntag_CRISPR/+* embryos labelled with *suntag* smFISH probes (see methods). Centered black bar represents the median, dashed black lines represent quartiles. ∗∗∗∗ p < 0.0001 with a two-tailed Welch’s t-test.

Remarkably, similar clustering of mRNA was also detected with the *twi_suntag* transgene (Figure S3e-f), where the mRNA distribution is different from that of the *twi_CRISPR a*lleles, owing to an initial erratic activation pattern (Figure S1f). Thus, mRNA clustering and co-translation do not seem to depend on mRNA concentration. To quantify these clusters and their ability to be translated, we divided the total pool of mRNAs into 4 distinct classes of mRNA densities with 1, 2, 3 or ≥4 mRNAs (Figure 3f) and scored their ability to undergo translation. Remarkably, molecules located in a large mRNA cluster were systematically engaged in translation. This is particularly true for mRNA clusters located at the basal perinuclear space (Figure 3f).

Given the enhanced clustering in this compartment, we questioned the properties of this basal compartment on translational efficiency. To estimate the efficiency of translation, we extracted the intensity of the scFv-GFP signal overlapping each individual mRNA molecule (not contained within a cluster) (see methods). We found that in the basal perinuclear space, a single molecule of mRNA is on average 50% more intense than a single molecule located apically, suggesting an enhanced efficiency of translation (Figure 3g). Collectively our results suggest that the basal perinuclear cytoplasm fosters an enhanced *twi* translation via the formation of co-translated mRNA clusters and via an enhanced efficiency of single mRNA molecules translation.

### Tracking of translating mRNPs reveals distinct mobilities

To quantitatively characterize the diffusive properties of *twi_suntag_CRISPR* mRNPs, we imaged, the scFv-GFP signal in the apical and basal perinuclear region in the mesoderm at high speed and employed single particle tracking (Figure 4a and movie S5, S6). The trajectories show clear differences in particle mobility between the apical and basal compartments (Figure 4b-c and Figure S4). The mRNPs located above nuclei tend to displace faster than those located below as quantified by 1D displacement without preferential drift (Figure S4b). This difference in behavior could be due to a differential confinement or alternatively and non-exclusively to a lower diffusion capacity of the mRNPs located at the basal perinuclear space. To discriminate between these two scenarios, we quantified the mean square displacement (MSD). MSD curves reveal clear distinct displacement properties among these two subpopulations (Figure 4d). In both populations, the sublinear growth of the MSD curves suggests a sub-diffusive behavior (also called anomalous diffusion)(*28*). Indeed, a linear fitting of the MSD curves was unsatisfactory (R^2^<0.8) (Figure S4 c). We therefore used a non-linear model (anomalous fitting)(*28, 29*) to fit to our data and estimated two parameters: the anomalous diffusion exponent (alpha parameter) and the diffusion coefficient. The anomalous alpha parameter was estimated to be <1 (Figure 4e), confirming the non-linearity and the confinement of the diffusion process. However, this parameter appears comparable in both compartments. Thus, the differential mobility of mRNPs between the apical and basal cytoplasmic compartments cannot be solely attributed to a distinctive confinement. We therefore examined the diffusion coefficient. Sites of translation located apically show an average diffusion coefficient of 0.051µm^2^ s^-1^, a value comparable to what has been reported in cell culture experiments using Suntag(*13, 18*) or by spt-PALM on ribosomes(*30*). Remarkably, the diffusion coefficient of mRNPs located basally is three-times lower, showing a significantly lower mobility (Figure 4f).

**Figure4:**
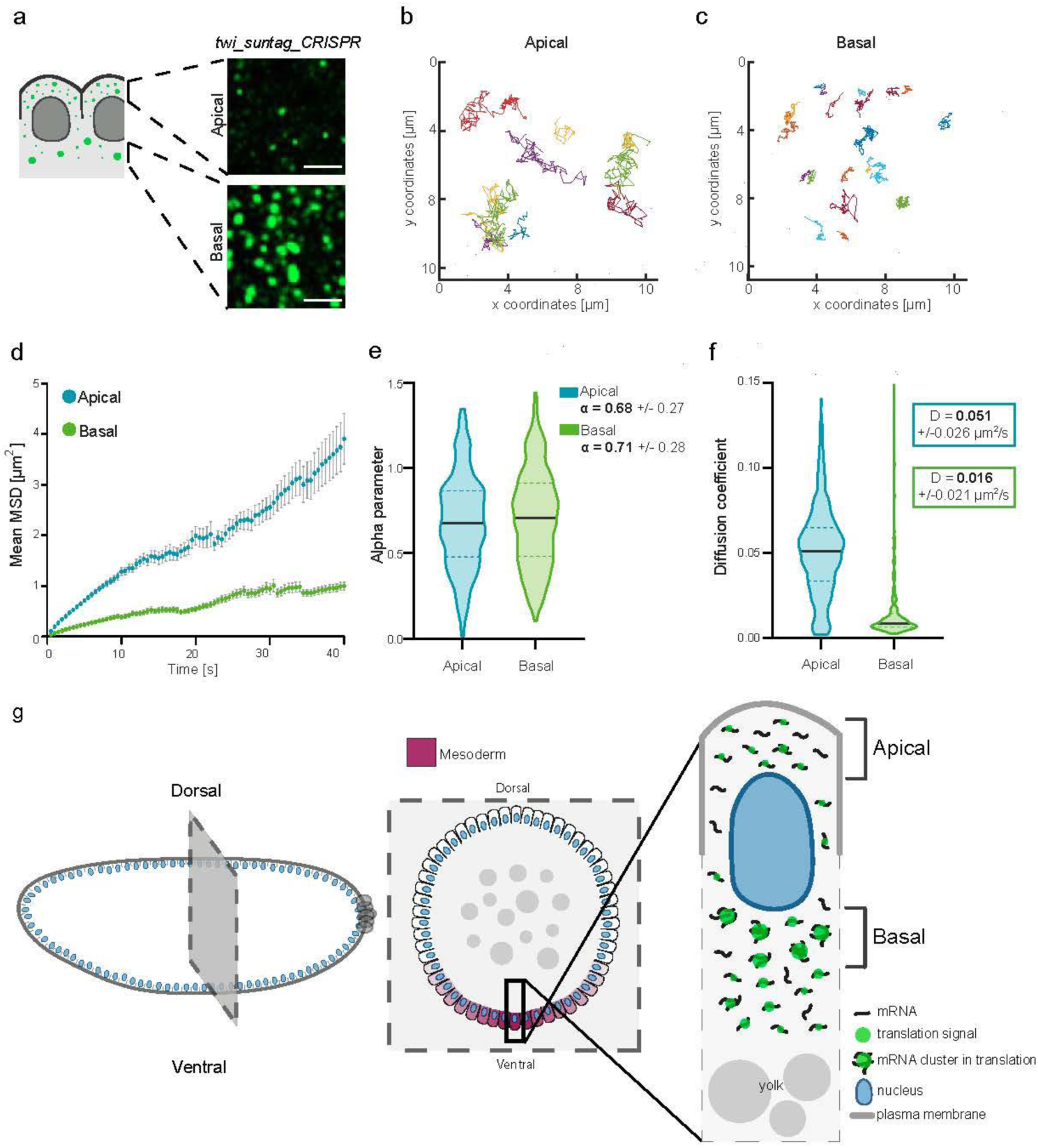
Quantifying the mobilities of *twist* mRNPs translation foci. **a**, Typical confocal images of translation foci located in theapical and basal perinuclear cytoplasm at nc14 used for single particle tracking. Images extracted from a fast movie at a rate of 2 frames s^-1^ of a living *scFv-GFP-NLS/+ > twi_suntag_CRISPR/+*embryo (related movie S5-6). Scale bars 5μm. **b, c**, Examples of color-coded trajectories of translation foci, taken from fast movies as shown in **a. d**, Graph representing the mean Mean Square Displacement (MSD) as a function of time for particles located apically (n=237 traces, three movies; blue) or basally (n=263 traces, three movies; green) nuclei. Error bars represent SEM. **e**, Violin plots of the alpha parameter for the particles located apically (blue) and basally (green) using anomalous fitting (centered black bar represent median, dashed lines represent quartiles). **f**, Violin plots of the estimated diffusion coefficient distribution of the particles located apically and basally of living *scFv-GFP-NLS/+ > twi_suntag_CRISPR/+* nc14 embryos (centered black bar represents median, dashed lines represent quartiles). **g**, Scheme of a nc14 embryo prior to cellularization, a corresponding sagittal section and a zoomed view of a pseudo-cell summarizing the spatial heterogeneity in *twist* translation.

In conclusion, these kinetic measurements of live translation spots (Figure 4) are consistent with the conclusions obtained from fixed tissues (Figure 3) and strongly suggest that in the basal perinuclear cytoplasm, translation sites primarily consist of slow diffusing clusters of mRNAs engaged in translation, referred to as translation factories. Low mobility of translation factories can be attributed to their larger size. However, an anchoring to particular organelles, specific to the basal perinuclear cytoplasm, could also slow down the diffusion of mRNPs and remains to be demonstrated.

## Conclusion

The Suntag method provides a versatile approach to measure translation kinetics of single mRNAs in a developing embryo. This labeling technology can be combined with mRNA labeling methods for the simultaneous quantitative imaging of transcription and translation dynamics. By focusing on *twi* mRNAs as a paradigm for transcription factor encoding transcripts, we have uncovered fundamental features of translation in a living organism. First, we have shown the existence of translation factories in an embryo, whereby clusters of several mRNAs were co-translated in a microenvironment, echoing what has recently been shown in cultured human cells(*18*) or in dividing yeast cells(*31*). Given the widespread subcellular localization of mRNA during development(*9*), we expect that our newly identified twi translation factories represent one example among many more to come. Co-translating multiple mRNAs within a local microenvironment may allow a higher efficiency *e.g* via ribosome recycling(*32, 33*). Secondly, *Drosophila twi* translation factories are not randomly distributed but rather concentrate in the basal perinuclear space. We discovered that in this compartment, the efficiency of translation of single mRNA molecules was significantly higher than that of identical mRNAs located elsewhere. This spatial bias might be supported by the already reported higher availability of mitochondria in this perinuclear compartment(*34*), in line with the fact that protein synthesis is one of the most energy-consuming processes in the cell(*35-37*).

In the context of a syncytial embryo, compartmentalization of the cytoplasm could be important for the confinement of protein synthesis and subsequent delivery to final functional destinations. In the case of *twi* mRNAs, which encodes a critical transcription factor, a combination of slow mRNPs mobility and enhanced efficiency of translation in the basal perinuclear space could limit the protein’s diffusion outside of the presumptive mesoderm and favor rapid import of neo-synthesized Twi protein to the nucleus. Rapid nuclear availability of this key transcription factor might be essential for rapid activation of the hundreds of direct target genes instructing the mesodermal fate(*27*). Enhancement of translation efficiency in dedicated cytoplasmic microenvironments could be employed by other developmental genes to ensure gene expression precision. Thus, precision in the establishment of developmental patterns(*38*), such as the presumptive mesoderm, cannot only be attributed to a precision in transcriptional activation. Future work will elucidate if other developmentally regulated mRNAs are translated via translation factories. We anticipate that the Suntag labeling method, combined with recently developed optogenetic manipulations(*39, 40*), will pave the way towards hitherto inaccessible translation modalities in intact developing organisms, much like the revolution triggered by the deployment of mRNA detection in living organisms.

## Methods

### *Drosophila* stocks and genetics

The *yw* stock was used as a control. *His2av-mRFP* (Bl23651) stock comes from Bloomington.*MCP-eGFP-His2Av-mRFP* comes from(*41*). For most experiments, female virgins expressing *scFv-sfGFP-NLS* were crossed to *yw* to obtain *scFv-sfGFP-NLS/+* flies. *scFv-sfGFP-NLS/+* virgins were crossed with males containing suntag repeats and/or MS2 constructs.

### Cloning and Transgenesis

The *twi-suntag* transgene was synthesized (GenScript Biotech) (Supplementary sequences 1) with 32x suntag repeats(*18*) into pUC57-simple. The *twi_suntag_MS2* transgene was generated based on the *twi_suntag* transgene with 128 MS2 repeats(*22*) inserted in the *XbaI* restriction site. Constructs were inserted into pbPHi(*42*) using *PmeI* and *FseI* and injected into BL9750 using PhiC31 targeted insertion^39^ (BestGene, Inc.). All *ScFv-sfGFP* lines were generated using NEBuilder® HiFi DNA Assembly Master Mix with primers listed in supplementary table 1 and inserted into pNosPE_MCP-eGFP (Supplementary sequences 1) after removal of MCP-eGFP. The recombination templates for CRISPR/Cas9 editing of *twist* gene to generate *twi_suntag_CRISPR* and *twi_suntag_MS2_CRISPR* were assembled with NEBuilder® HiFi DNA Assembly Master Mix (primers listed in supplementary table 1) and inserted into pBluescript opened *KpnI* and *SacI*. The *twi-suntag* transgene was digested with *FseI* and *SacII* and inserted into the recombination template opened with *FseI* and *SacII*, 128x MS2 repeats(*22*) were inserted in an *XbaI* restriction site. Guide RNA (supplementary table 1) were cloned into pCFD3-dU6:3gRNA (Addgene 49410) digested by *BbsI* using annealed oligonucleotides. Recombination template and guide RNAs were injected into BDSC#55821 (BestGene Inc.) and transformant flies were screened using dsRed marker inserted after the 3’UTR of *twi_suntag_CRISPR* and *twi_suntag_MS2_CRISPR. MCP-TagRFPT* constructs were assembled by replacing the eGFP fragment of pNosPE_MCP-eGFP using *NheI/BamHI* with the TagRFPT coding sequence amplified by PCR (supplementary table 1) from TagRFP-T-Rabenosyn-5 (Addgene 37537). *MCP-TagRFPT-NLS* was generated by insertion of the TagRFPT-NLS coding sequence into pNosPE_MCP-eGFP with NEBuilder® HiFi DNA Assembly Master Mix (primers listed in supplementary table 1).

### Single molecule fluorescence in situ hybridization (smFISH) and Immunostaining

Embryos were dechorionated with bleach for 3 min and thoroughly rinsed with H2O. They were fixed in a 1:1 solution of 10% formaldehyde:100% heptane for 25 min with shaking. Formaldehyde was replaced by methanol and embryos were vortexed for 1min. Embryos that sank to the bottom of the tube were rinsed three times with methanol. For immunostaining, embryos were rinsed with methanol and washed three times with PBT (PBS 1× 0.1% Triton X-100). Embryos were incubated on a rotating wheel at room temperature twice for 30 min in PBT, once for 20 min in PBT+ 1% BSA, and at 4 °C overnight in PBT 1% BSA with guinea-pig anti-Twist 1/200 (gift from Robert Zinzen). Embryos were rinsed three times and washed twice for 30 min in PBT, then incubated in PBT+ 1% BSA for 30 min, and in PBT +1% BSA with antibodies anti-guinea-pig Alexa 555-conjugated (Life technologies, A21435) 1/500 for 2 h at room temperature. Embryos were rinsed three times then washed three times in PBT for 10 min. DNA staining was performed using DAPI at 0.5 μg.ml^−1^.

Single molecule Fluorescent *in situ* hybridization was performed as follows: wash 5min in 1:1 methanol:ethanol, rinse twice with ethanol 100%, wash 5min twice in ethanol 100%, rinse twice in methanol, wash 5min once in methanol, rinse twice in PBT-RNa (PBS 1×, 0.1% tween, RNasin® Ribonuclease Inhibitors). Then, embryos were washed 4 times for 15 min in PBT-RNa supplemented with 0.5% ultrapure BSA and then once 20 min in Wash Buffer (10% 20X SCC, 10% Formamide). They were then incubated overnight at 37°C in Hybridization Buffer (10% Formamide, 10% 20X SSC, 400µg/ml tRNA, 5% dextran sulfate, 1% vanadyl ribonucleoside complex (VRC) and anti-suntag Stellaris probes coupled to Quasar 570 or Quasar 670 and/or anti-twist Stellaris probes coupled to Quasar 570. Probe sequences are listed in supplementary table 1. Probes against 32X MS2 coupled to Cy3 are a kind gift from Edouard Bertrand. Embryos were washed in Wash Buffer at 37°C and then in 2X SCC, 0.1% Tween at room temperature before mounting (with Pro-Long® Diamond antifade reagent). Images were acquired using a Zeiss LSM880 confocal microscope with a Airyscan detector in SR mode with a 40x Plan-Apochromat (1.3NA) oil objective lens, a 63X Plan-Apochromat (1.4NA) oil objective lens or a 20x Plan-Apochromat (0.8NA) air objective lens. GFP were excited using a 488nm laser, Cy3 and Quasar570 were excited using a 561nm laser, Quasar670 was excited using a 633nm laser.

### Image analysis

Analysis of smFISH data related to figure 3 was performed by custom-made algorithms developed in Python™. Briefly, a blob detection was performed on the scFv-GFP and mRNA channels (green and red respectively) separately. Raw data were filtered frame by frame with a two-dimensional Difference of Gaussian Filter whose kernels size are determined as in(*38*) and the filtered images were thresholded with a user defined threshold value; the choice of the threshold was driven by visual inspection thanks to an interactive graphical tool. All the 3D connected components of the resulting binary images are considered as spots, which are then filtered in size with a volume threshold of 10 pixels. All spots with centroids in the same frame where the nuclei were detected were removed so as to only analyze mRNAs located at the apical and basal perinuclear cytoplasmic part. Within the detected mRNA pool, three populations could be distinguished by volume or intensity. As expected, smFISH allowed us to detect single molecules of mRNA, mRNAs in clusters and mRNAs undergoing decay. By looking at the mRNA spots not engaged in translation (red spots not overlapping green spots), a separable population can be characterized by a very low spot volume. We speculate that these small spots are mRNA in decay that are unable to be translated. To remove these small mRNA spots (possibly in decay), we refined the volume threshold of a single molecule mRNA by looking at the histogram of the volume of the mRNA spots in translation (red spots overlapping with at least one pixel of a green spot). This led to the detection of two major populations on the histogram: single molecules of mRNA, representing the larger population shown in the histogram, and mRNA clusters, characterized by a smaller population and a bigger volume. We performed a double Gaussian fitting on the histogram to differentiate the single molecule and cluster populations, then we defined a threshold value for the lower bound of the single molecule volume as µ – 3σ, where µ and σ are, respectively, the mean and the standard deviation of the Gaussian function fitting single molecule mRNA spots. Having obtained the correct threshold value for the volume, we reanalyzed the mRNA channel to connect the spots detection results from both channels. We obtained two clear populations of mRNAs that we divided into two subpopulations based on position above or below the nucleus.

The first parameter extracted is the percentage of mRNA in translation: given by the ratio between the number of mRNA spots overlapping a translation spot, divided by the total number of mRNA spots.

We also extracted the efficiency of translation. For this, we estimated mRNA single molecule intensity separately for above and below the nucleus by performing a Gaussian fitting on the histogram of intensities of all detected mRNA spots. Single molecules are by far the widest population, so a single Gaussian fitting gives an estimate of the single molecule intensity, not influenced by the population of mRNA clusters, also present in the histogram. We used µ + 3σ for the single molecule upper intensity bound where µ and σ are the mean and the standard deviation of the fitting respectively.

For translation spots we used the GFP intensity divided by background, where the background is calculated for each spot as the average intensity value of the pixels surrounding the spot itself. This allowed us to rescale by the background, because in this case we have a bath of free diffusing GFP (provided by the *nos-scFv-GFP-NLS* transgene).

To score differential clustering of mRNAs, above and below nuclei, all mRNA spots intensities above µ + 3σ (mRNA clusters) were rescaled by the mean intensity of a single mRNA molecule (extracted from the Gaussian fitting), which allowed us to express the signal of each mRNA entity (cluster of varying sizes) as a function of a single molecule. For translation efficiency and clustering of mRNA, outliers were removed using the ROUT method(*43*), embedded in GraphPad Prism 8 software, with Q set to 0.1%.

We then classified all the spots in terms of intensity, assigning a value of 1 to a spot if its intensity is < µ + 3σ, value 2 if its intensity I, is such that µ + 3σ < I < 2 * (µ + 3σ) and so on. For each spot category, we checked the percentage of mRNA spots overlapping a translational spot, meaning that they are engaged in translation. For smFISH of the CRISPR alleles, analysis above the nucleus contained between 5 and 8 confocal Z-planes spaced 0.5μm apart, and analysis of the region below the nucleus contained 17 to 28 Z-planes spaced 0.5μm apart. For smFISH analysis with *twi_suntag* transgene, only one filtering with a volume threshold of 10 pixels was performed as only two populations appeared (single molecule mRNA and mRNA clusters). Analysis for above the nucleus contained 6 to 12 confocal Z-planes spaced 0.5μm apart and analysis below the nucleus contain 36 to 41 confocal Z-planes spaced 0.5μm apart.

### Light-Sheet Microscopy

For light-sheet imaging (related to movie 2,5,6 and S3-4), we employed the MuViSPIM (Luxendo, a Brüker company). This setup provides two-sided illumination with two Nikon 10x/0.3 water objectives and two-sided detection with two Olympus 20x/1.0 W objectives. The light sheet is generated by a scanning of a gaussian beam. We used the line illumination tool for improved background suppression. Images are acquired by two ORCA Flash 4.0 (C91440) from Hamamatsu and processed by LuxControl v1.10.2. A 50ms exposure time was used for the green and red channel with a 488nm and 561nm laser excitation respectively. Maximum intensity projection were processed with Fiji(*44*). Fusion is processed by a software solution from Luxendo (Image Processor v2.9.0) Deconvolution was performed after the fusion process and executed with Huygens Professional v 19.10 (Scientific Volume Imaging B.V). We used the gaussian multi-view light sheet parameters for processing of 3D+t images. 3D reconstruction was done with Imaris v9.5.0 (Bitplane, an Oxford company). The ortho slicer tool was used to show cross sections of embryos with ≈5µm extended section thickness for movie 6 and S4 and ≈10µm extended section thickness for movie S3.

### Live Imaging

Embryos were dechorionated with tape and mounted between a hydrophobic membrane and a coverslip as described previously^14^. Movies for *scFv-GFP*-noNLS (related to movie S1) and *scFv-GFP-NLS* (related to movie 1) were acquired using a Zeiss LSM780 confocal microscope with a Plan-Apochromat 40x/1.3 oil objective lens, with GFP excitation using a 488nm laser. A GaAsP detector was used to detect GFP fluorescence with the following settings: 1024× 1024 pixels, each Z-stacks comprised 6 planes, 16-bit and zoom 2.0. Movies for scFv-GFP-Bcd3’UTR-NLS were acquired using a Zeiss LSM780 confocal microscope with a Plan-Apochromat 40x/1.3 Oil objective and the following settings: 512× 512 pixels, each Z-stacks comprised 12 planes, 16-bit. Movies of *scFv-GFP-NLS/+*> *twi_suntag_CRISPR/+*or *twi_suntag_MS2_CRISPR/+* were acquired using a Zeiss LSM880 with confocal microscope in fast Airyscan mode with a Plan-Apochromat 40x/1.3 oil objective lens. GFP was excited using a 488nm laser with the following settings: 640× 640 pixel images, each Z-stacks comprised 50-60 planes spaced 0.5μm apart and zoom 2.5x. Movies were then processed to remove frame outside of the embryos or containing the membrane signal to correct drifting, and processed stacks were max intensity projected using a custom made software, developed in Python™. Movies of *scFv-GFP-NLS/MCP-TagRFPT-NLS* >*twi_suntag_MS2/+ transgene* (related movie 4) were acquired using a Zeiss LSM880 with confocal microscope in fast Airyscan mode with a Plan-Apochromat 40x/1.3 oil objective lens. GFP and RFP were excited using a 488nm and 561nm laser respectively with the following settings: 276× 316 pixel images, each Z-stacks comprised 2 planes spaced 0.5μm apart and zoom 14x.

### Puromycin Injection

Embryos were dechorionated with tape, lined up on a hydrophobic membrane covered in heptane glue with desiccation at room temperature for 10 minutes prior to being covered in Voltalef Oil 10S (VWR), and injected with 10mg/mL puromycin (InvivoGen) using a FemtoJet ***5427*** (Eppendorf) micro-injector and associated Femtotips® I (Eppendorf) needles. Embryos were injected in the lateral region immediately prior to coverslip positioning. Time lapse images (related to Movie 3) were acquired using a confocal LSM780 (Zeiss) microscope. A GaAsP detector was used to detect the GFP fluorescence excited using a 488nm laser.

### Detection and tracking of single particles

For single particle tracking (related movie S5-6), movies were acquired with a Zeiss LSM880 with confocal microscope in fast Airyscan mode with a Plan-Apochromat 40x/1.3 oil objective lens, GFP was excited using a 488nm laser and the following settings: 132 × 132 pixels, each Z-stacks comprised 10 planes 0.5μm apart, Zoom 8x. Under these conditions a z-stack was acquired every ≈500ms.

Each movie was max intensity Z-projected using Fiji(*44*). Single particle trajectories were generated with MatLab 2015b (Mathworks Inc., USA) using SLIMfast, which implements the Multiple-Target-Tracing algorithm(*45, 46*). Individual spots were localized using a blob diameter of 9 pixels. They were then tracked using maximal frame gap of 0 frame and a maximal expected diffusion coefficient set to D_max_ = 0,5 µm^2^.s^-1^. Tracked movies were then evaluated using the script evalSPT(*47*). Each tracked spots was visually checked and poorly tracked spots were manually removed using evalSPT. In total we obtained 237 traces from three movies above nuclei (Apical) and 263 traces from three movies below nuclei (Basal).1D displacement in x and y were obtained from evalSPT tool for each movie. Mean square displacements (MSD) were calculated for tracks present for at least 10 consecutive frames using the MSDanalyzer MatLab script(*48*).

Coefficients diffusion were first obtained by fitting 0 to 20 seconds of the MSD of individual trajectories with a linear model as follows:

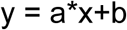

To estimate MSD, according to Einstein’s theory, the MSD of Brownian motion is described as

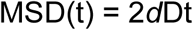

with *d* representing dimensionality, in our case, *d* = 2. D represents the diffusion coefficient.

Then, MSD of individual trajectories were fitted with a non-linear model, as follows:

If x = log(D) and y = log(MSD) where ‘D’ are the delays at which the MSD is calculated, then this method fits y = f(x) by a straight line

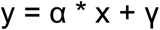

so that we approximate the MSD curves by MSD = γ * D^α^.

According to a previous modeling study(*49*), an α value of 1 indicates free diffusive movement, a value of α close to 2 indicates movement by active transport, and a value of α <1 indicates a motion constrained in space.

## Acknowledgements

We are grateful to E.Bertrand and X.Pichon for sharing the 32x suntag plasmid and MS2-Cy3 smFISH probes and helpful discussions. We thank I.Izzedin for providing evalSPT and MSD analyser Matlab scripts. We thank R.Zinzen for sharing the twi antibody. We thank C.Desplan, R.Bordonne, T.Saunders, V.Pimmett, and all members of the Lagha lab for their critical reading of the manuscript and constructive discussions. We acknowledge the MRI imaging facility, a member of the national infrastructure France-BioImaging supported by the French National Research Agency (ANR-10-INBS-04, «Investments for the future»). M.B is a recipient of an FRM fellowship. This work was supported by the ERC SyncDev starting grant to M.L and a HFSP-CDA grant to M.L. M.L, J.D and S.D.R are sponsored by the CNRS.

## Author Contribution

M.L conceived the project. M.L and J.D designed the experiments. J.D, M.B and M.D performed experiments. A.T developed software for image and data analysis. M.B and J.D performed kinetics diffusion analysis. S.D.R and J.D performed MuViSPIM imaging and image processing. J.D, M.L, A.T, and M.B analyzed the data. J.D, M.L, and M.B interpreted the results. M.B performed all artwork. M.L wrote the manuscript with help from J.D. All authors discussed, approved and reviewed the manuscript

## Supplementary figure

**Figure S1:**
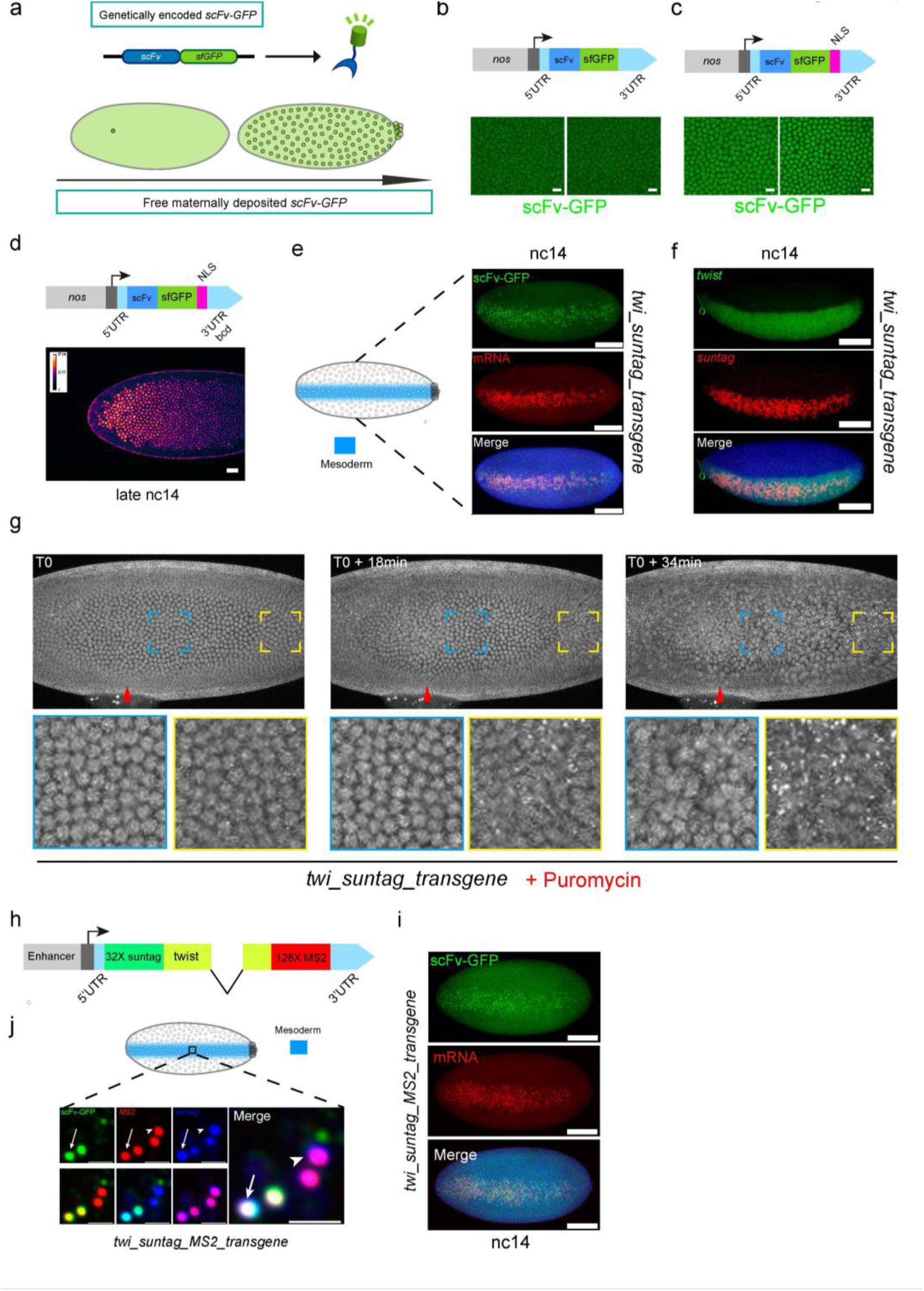
Imaging translation with suntag labeling in *Drosophila* embryos. **a**, Schematic representation of the genetically encoded and maternally deposited scFv-GFP detector lines created in this study. **b, c**, Snapshots of maximum intensity projection of confocal time lapse movie of *Drosophila* embryos expressing the *scFv-GFP* (**b**) or the *scFv-GFP-NLS* (**c**) transgene during the nc14. NLS: nuclear localization signal, scale bars 10μm. Related movies S1 and 1 respectively. **d**, Heatmap of GFP fluorescence at late nc14, showing an enhanced fluorescence at the anterior part of the *scFv-GFP-NLS-bcd3’UTR* transgenic embryo, scale bars 20μm. **e**, Maximum intensity projection of a ventral view of a *scFv-GFP-NLS/+ > twi_suntag/+* transgenic embryo in nc14 stained with endogenous scFv-GFP (green) and smFISH probes against suntag (red), scale bars 100μm. **f**, Maximum intensity projection of confocal images from a nc14 *scFv-GFP-NLS/+ >twi_suntag/+* transgenic embryo, stained with probes against *twist* (green) and *suntag* repeats (red). Scale bars 100μm. **g**, Maximum intensity projection of frames taken from live imaging (related movie 3) of a *scFv-GFP-NLS/+ > twi_suntag/+* embryo injected with puromycin. Red arrowheads indicate the site of drug injection. T0 correspond to ≈25-30min after puromycin injection. Zoomed images from the color-coded indicated regions are provided in the lower panels. **h**, Schematic of the *twi_suntag_MS2* transgene. **i**, Maximum intensity projection of confocal images from a nc14 *scFv-GFP-NLS/+ >twi_suntag_MS2/+* transgenic embryo, stained with endogenous scFv-GFP (green) and smFISH probes against suntag repeats (red). Scale bars 100μm. **j**, Zoomed confocal plane of a *scFv-GFP-NLS/+ > twi_suntag_MS2/+* transgenic embryo stained with endogenous scFv-GFP (green) and probes against *suntag* (blue) and *MS2* (red) repeats. scFv-GFP foci colocalizing with *suntag* and *MS2* probes reveal mRNAs in translation (arrow), while spots that are only stained with *suntag* and *MS2* probes represent mRNAs not in translation (arrowhead), scale bars 1μm.

**Figure S2:**
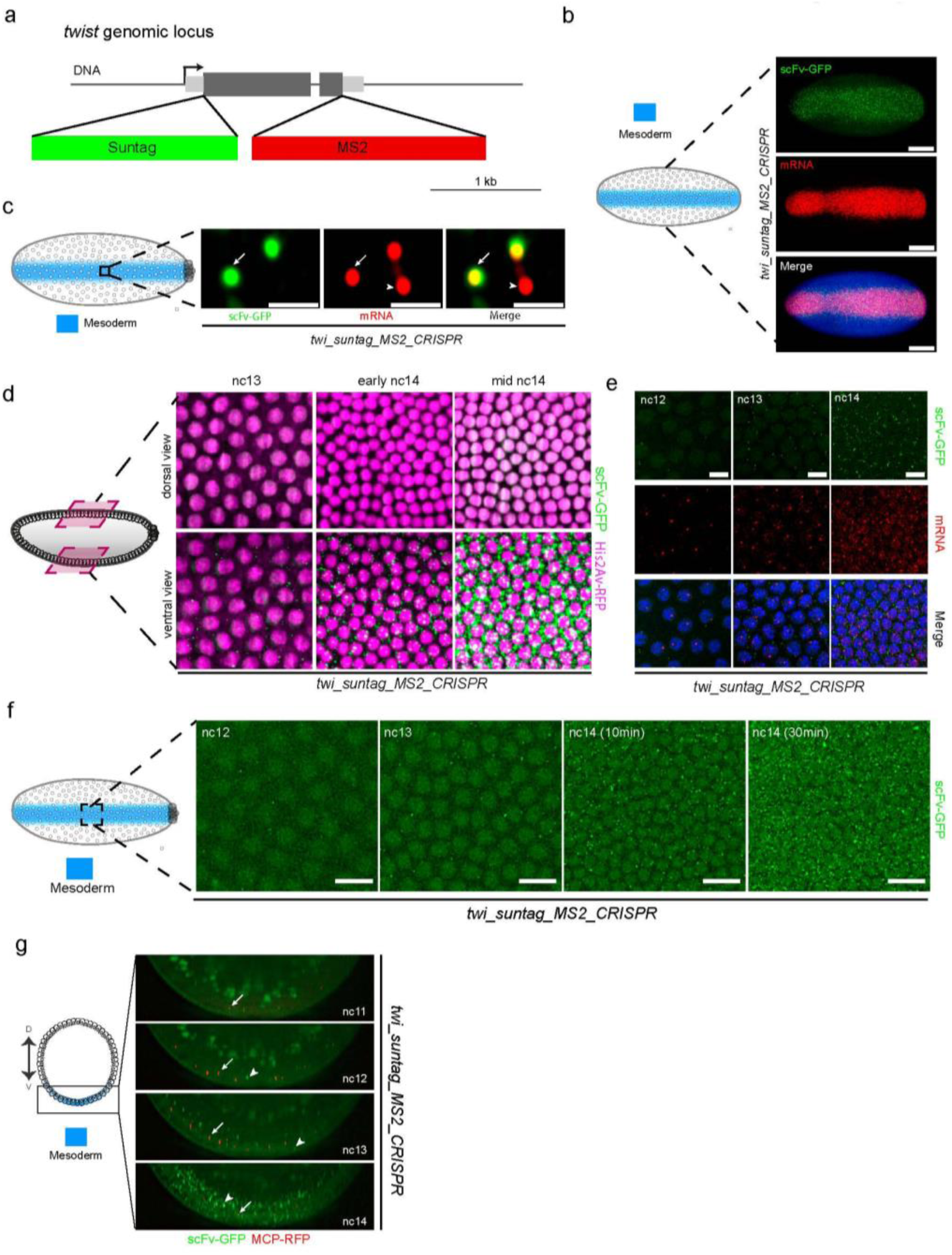
Capturing the timing of translation of endogenous *twist* mRNAs. **a**, Schematic of the *twi_suntag_MS2_CRISPR* targeting strategy. **b, c**, Maximum intensity projection of confocal images of nc14 *twi_suntag_MS2_CRISPR Drosophila* embryos expressing scFv-GFP (green) labelled with *suntag* probes (red), scale bars 100μm. **c**, Zoomed view within the mesoderm shows two distinct mRNA (red) pools: those engaged in translation (co-localizing with scFv-GFP signal,arrows)and untranslated single molecules red, arrowheads). Scale bars 1μm. **d**, Maximum intensity projection snapshots of *His2Av-mRFP/+; scFv-GFP-NLS/+ > twi_suntag_MS2_CRISPR/+* embryo imaged with a MuVi-SPIM, (related movie S2). Upper panels show the dorsal part of the embryo while the lower panels show the ventral mesoderm part of the embryo, where *twi* is expressed. Nuclei are shown in magenta and sites of translation in green. **e**, Maximum intensity projection of confocal images from nc12, nc13 and nc14 *twi_suntag_MS2_CRISPR* embryos expressing scFv-GFP (green) and labelled with *suntag* probes (red) and DAPI (blue). Scale bar, 10μm. **f**, Maximum intensity projection of confocal images of a *scFv-GFP-NLS/+ > twi_suntag_MS2_CRISPR/+* live embryo imaged from nc12 to nc14 showing *twi* translational activation. Scale bar*+*s 10μm. **g**, Sagittal view of a confocal time lapse movie of a *scFv-GFP-NLS/MCP-TagRFPT >Twi_suntag_MS2_CRISPR/+* embryo (related movie S3). Sites of nascent transcription appear as bright red spots (arrows) and sites of translation as green spots (arrowheads). Left panel: schematic of a sagittal section throughout an embryo, with the dorso-ventral axis (D, V) and the presumptive mesoderm (blue cells) indicated.

**Figure S3:**
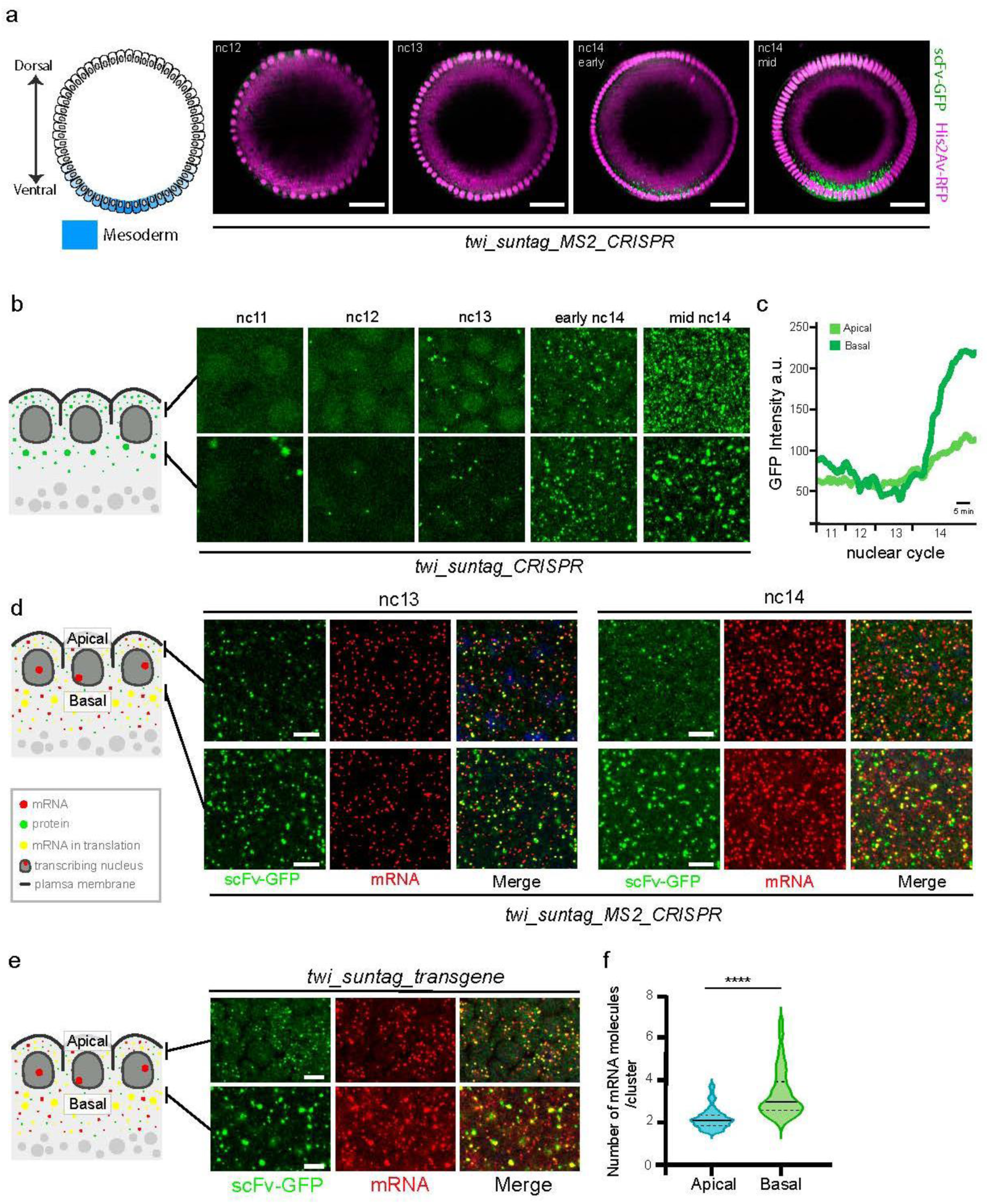
Revealing a spatial heterogeneity in translation sites in the mesoderm. **a**, Representative cross section of a transverse plane from reconstructed MuViSPIM images of a *His2Av-mRFP/+; scFv-GFP-NLS/+ > twi_suntag_MS2_CRISPR/+* embryo (movie S4). Ventral mesodermal nuclei are located at the bottom. Nuclei are observed with a *His2Av-mRFP* transgene (magenta) and sites of translation with the scFV-GFP (green dots). Scale bars 30μm. **b**, Confocal images of a *scFv-GFP-NLS/+ > twi_suntag_CRISPR/+* live embryo imaged from nc11 to nc14. Upper panels show a maximum intensity projection of four z-stacks located apically (≈2 μm). Lower panels show a maximum intensity projection of 21 z-stacks located basally (≈10 μm). **c**, Quantification of total intensities of GFP foci apically (light green) and basally (dark green) shown in panel b. A schematic representation of the two compartments analyzed is provided on the left. **d**, Maximum intensity projection of three z-stacks (≈1 μm) of confocal images of nc13 and nc14 *scFv-GFP-NLS/+ > twi_suntag_MS2_CRISPR/+ Drosophila* embryos expressing scFv-GFP (green) labelled with *suntag* probes (red) and DAPI (blue), scale bars 5μm. **e**, Maximum intensity projection of three z-stacks (≈1 μm) of confocal images of nc14 *scFv-GFP-NLS/+ > twi_suntag/+* transgenic *Drosophila* embryos expressing scFv-GFP (green) labelled with *suntag* probes (red) and DAPI (blue), scale bars 5μm. **f**, Violin plot of the distribution of the number of mRNA molecules per cluster quantified from images taken apically (n=207; blue) and basally (n=688; green) from three nc14 embryos *scFv-GFP-NLS/+ > twi_suntag/+* transgene labelled with *suntag* smFISH probes. Centered black bar represent median, dashed black lines represent quartiles. ∗∗∗∗ p < 0.0001 with a two-tailed Welch’s t-test.

**Figure S4:**
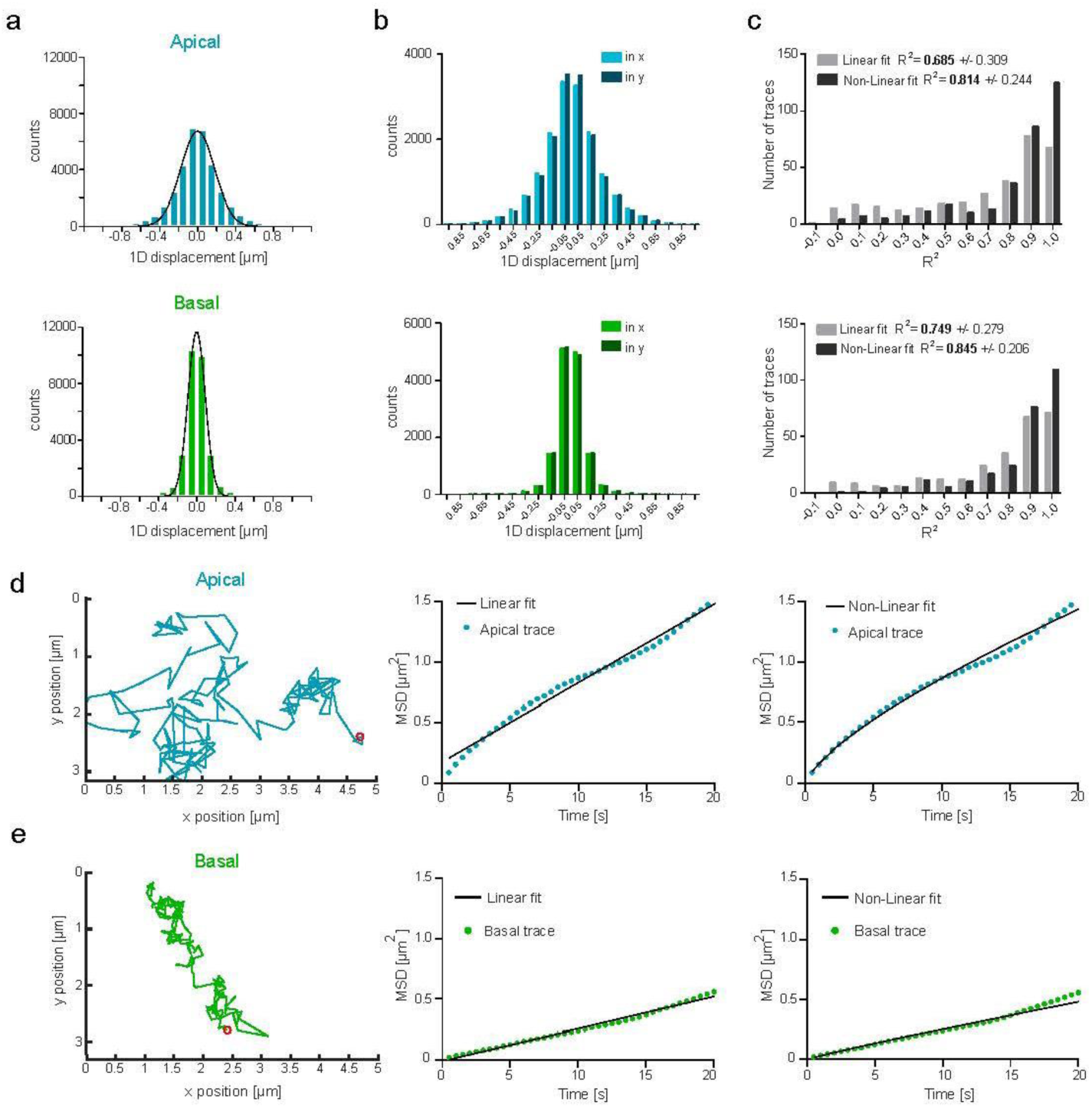
Quantifying the mobilities of *twi* mRNAs translation sites. **a**, 1D displacement distribution in x and y merged measured between two consecutive frames of traces located in the apical (blue) and basal (green) perinuclear cytoplasm of *scFv-GFP-NLS/+ > twi_suntag_CRISPR/+* embryos. Gaussian fitting is shown in black. **b**, 1D displacement distribution in x and in y separately measured between two consecutive frames of traces located in the apical perinuclear cytoplasm (top histogram, x (light blue) and y (dark blue)) and basal perinuclear cytoplasm (bottom histogram, x (light green) and y (dark green)). Displacement are equally distributed in x and y showing no drift due to cellular movements. **c**, Distribution of R^2^ as a goodness of the linear fit (light grey) and non-linear fit (dark grey) of apical traces (top histogram) and basal traces (bottom histogram). A good fit is considered when R^2^>0.8. **d**, One representative trajectory of a translation particle located apically (left panel) as well as linear (middle panel) and non-linear (right panel) fitting of its MSD over time. **e**, One representative trajectory of a translation particle located basally (left panel) as well as linear (middle panel) and non-linear (left panel) fitting of its MSD over time.

## List of Movies and Movies Legend

**Movie 1**: Maximum intensity projection of confocal Z-stacks live imaging of an embryo containing scFv-sfGFP-NLS showing no formation of aggregates, scale bars 10μm.

**Movie 2**: Maximum intensity projection of LightSheet Z-stacks live imaging of a *scFv-sfGFP-NLS/+ > twi_suntag/+* transgenic embryo.

**Movie 3**: Maximum intensity projection of confocal Z-stacks live imaging of *scFv-sfGFP-NLS/+ > twi_suntag* transgenic embryo after Puromycine injection, scale bars 50μm.

**Movie 4**: Maximum intensity projection of 2z plane confocal live imaging of *scFv-sfGFP-NLS/MCP-TagRFPT-NLS > twi_suntag/+* transgenic embryo, scale bars 1μm.

**Movie 5**: Maximum intensity projection of LightSheet Z-stack live imaging of *His2Av-mRFP*/+; *scFv-sfGFP-NLS/+ > twi_suntag_CRISPR/+* embryo. Upper part: dorsal view, lower part: ventral view. Nuclei are detected using His2Av-mRFP and suntag using scFv-GFP.

**Movie 6**: Reconstructed 5µm thick movie of a MuViSPIM live imaging of a *His2Av-mRFP/+; scFv-GFP-NLS/+ > twi_suntag_ CRISPR/+* embryo. Nuclei are detected using His2Av-mRFP and suntag using scFv-GFP.

## Supplementary Movies

**Movie S1**: Maximum intensity projection of confocal Z-stacks live imaging of an embryo containing scFv-sfGFP no NLS showing no formation of aggregates, scale bars 10μm.

**Movie S2**: Maximum intensity projection of Light-Sheet Z-stack live imaging of *His2Av-mRFP*/+; *scFv-sfGFP-NLS/+ > twi _suntag_MS2_CRISPR/+* embryo. Upper part: dorsal view, lower part: ventral view. Nuclei are detected using His2Av-mRFP and suntag using scFv-GFP.

**Movie S3:** Maximum intensity projection of LightSheet Z-stack 10µm thick live imaging of *scFv-sfGFP-NLS, MCP-TagRFPT/+ > twi _suntag_MS2_CRISPR/+* embryo. Transcription sites are detected using MCP-TagRFPT and suntag using scFv-GFP.

**Movie S4**: Reconstructed 5µm thick movie of a MuViSPIM live imaging of a *His2Av-mRFP/+; scFv-GFP-NLS/+ > twi_suntag_MS2_CRISPR/+* embryo. Nuclei are detected using His2Av-mRFP and suntag using scFv-GFP. **Movie S5**: Maximum intensity projection of confocal Z-stacks live imaging of an embryo containing *scFv-sfGFP-NLS/+ > twi_suntag_CRISPR*. Translation foci above nuclei are shown (imaging are at 2FPS, movie are shown at 12FPS), scale bars 5μm.

**Movie S6:** Maximum intensity projection of confocal Z-stacks live imaging of an embryo containing *scFv-sfGFP-NLS/+ > twi_suntag_CRISPR*. Translation foci below nuclei are shown (imaging are at 2FPS, movie are shown at 12FPS), scale bars 5μm.

